# Anesthesia-induced loss of consciousness disrupts auditory responses beyond primary cortex

**DOI:** 10.1101/502385

**Authors:** Aaron J Krom, Amit Marmelshtein, Hagar Gelbard-Sagiv, Ariel Tankus, Hanna Hayat, Daniel Hayat, Idit Matot, Ido Strauss, Firas Fahoum, Martin Soehle, Jan Boström, Florian Mormann, Itzhak Fried, Yuval Nir

## Abstract

Despite its ubiquitous use in medicine, and extensive knowledge of its molecular and cellular effects, how anesthesia induces loss of consciousness (LOC) and affects sensory processing remains poorly understood. Specifically, it is unclear whether anesthesia primarily disrupts thalamocortical relay or intercortical signaling. Here we recorded intracranial EEG (iEEG), local field potentials (LFPs), and single-unit activity in patients during wakefulness and light anesthesia. Propofol infusion was gradually increased while auditory stimuli were presented and patients responded to a target stimulus until they became unresponsive. We found widespread iEEG responses in association cortices during wakefulness, which were attenuated and restricted to auditory regions upon LOC. Neuronal spiking and LFP responses in primary auditory cortex (PAC) persisted after LOC, while responses in higher-order auditory regions were variable, with neuronal spiking largely attenuated. Gamma power induced by word stimuli increased after LOC while its frequency profile slowed, thus differing from local spiking activity. In summary, anesthesia-induced LOC disrupts auditory processing in association cortices while relatively sparing responses in PAC, opening new avenues for future research into mechanisms of LOC and the design of anesthetic monitoring devices.

## Introduction

The practice of administering general anesthesia, which began over 170 years ago, revolutionized medicine by transforming surgery from being a deeply traumatic experience into a humane therapy(1, 2) and enabling complex procedures to be performed. In the US, nearly 60,000 patients per day undergo general anesthesia(3): a drug-induced, reversible condition that includes unconsciousness, amnesia, analgesia, and akinesia. By now, we have extensive knowledge of the molecular and cellular effects of anesthetics(4–7); but how anesthetic drugs induce loss of consciousness (LOC) remains a central unresolved question in medicine(8). Given that anesthetics act at diverse molecular and cellular targets(7, 9), their effects on consciousness and sensory perception are best studied at the level of common brain circuits and systems(2, 9, 10). Indeed, recent studies focusing on the systems level effects of anesthesia proposed that LOC may involve sleep pathways(9, 11), thalamocortical circuits(12), specific brainstem regions(2, 13, 14), or unbinding of fronto-parietal cortical activities(15, 16).

Traditionally, sensory pathways were studied in anesthetized animals where robust responses in primary sensory regions prompted seminal discoveries on the organizational principles in multiple modalities(17–19). With time, research expanded to awake animals and a number of studies directly compared sensory processing across wakefulness and anesthesia, revealing state-dependent evoked responses in the visual(20–22) and somatosensory domains(23, 24). However, these modalities heavily depend on active sensing (i.e. eye movements, whisking)(25) and its absence in anesthesia limits interpretation as reflecting LOC. In the auditory domain where processing is largely passive, anesthesia was originally reported to reduce responses in auditory cortex(26–28), but recent studies with light anesthesia challenge this view(29). In humans, effects of anesthesia have been reported for mid-latency cortical evoked potentials(30) and for high-frequency (40Hz) steady-state potentials(31). Functional magnetic resonance imaging (fMRI) studies revealed differential responses across wakefulness and anesthesia(32–34), but interpretation is limited by poor temporal resolution and by relying on hemodynamics that change upon anesthesia(35). Generally, previous studies that compared sensory processing across wakefulness and anesthesia mostly focused on activity up to primary sensory cortex(26–28), often employed deep anesthesia at surgical levels(31, 34), entailed uncertainty about the precise moment of LOC in animal models, or relied on non-invasive measures in humans with limited resolution(31–34, 36). We sought to go beyond these limitations to better understand how anesthesia-induced LOC affects sensory processing. In particular, we aimed to address a central unresolved question in the field: whether anesthesia-induced LOC primarily affects bottom-up thalamocortical relay to primary sensory regions (‘thalamic gating’(37, 38)) or, alternatively, whether it mainly affects cortical signaling beyond primary regions(12).

We capitalized on a unique opportunity to compare auditory responses in neurosurgical epilepsy patients implanted with depth electrodes as they were anesthetized immediately before removal of intracranial electrodes. Extending protocols successfully employed to study anesthetic-LOC (39–41), we recorded intracranial field potentials and neuronal spiking activity to compare auditory responses in wakefulness and light “just-hypnotic” anesthesia, allowing to focus on changes related to LOC. Working with human subjects capable of responding purposefully to verbal tasks allowed us to reliably assess the moment of LOC. High-fidelity auditory responses evident in individual trials enabled comparison of sensory responses in short intervals immediately before and after LOC, thereby isolating the changes most relevant to LOC and minimizing confounding effects of deep anesthesia (e.g. changes in breathing, blood circulation, and thermoregulation). Our results reveal that anesthesia-induced LOC disrupts auditory responses in association cortex, while significant early responses in primary auditory cortex are relatively preserved. This distinction was observed for both simple (click-train) and complex (word) stimuli, and evident in LFPs as well as in single-neuron spiking activity representing neuronal output.

## Results

To study the effects of anesthesia-induced LOC on sensory-evoked neuronal activity in humans, we recorded intracranial EEG (iEEG), Local Field Potentials (LFPs), and neuronal spiking activity in epilepsy patients implanted with depth electrodes (Fig. 1a,b) or subdural cortical grids/strips, as they were anesthetized immediately before removal of intracranial electrodes. In nine experimental sessions (Supplementary Table 1), individuals listened to 40Hz click trains, and from the third session onwards, they were also presented with words and performed an auditory detection task. During each session the subjects were gradually anesthetized with propofol (Fig. 1c,d & Supplementary Figs. 1,2).

**Fig. 1.**
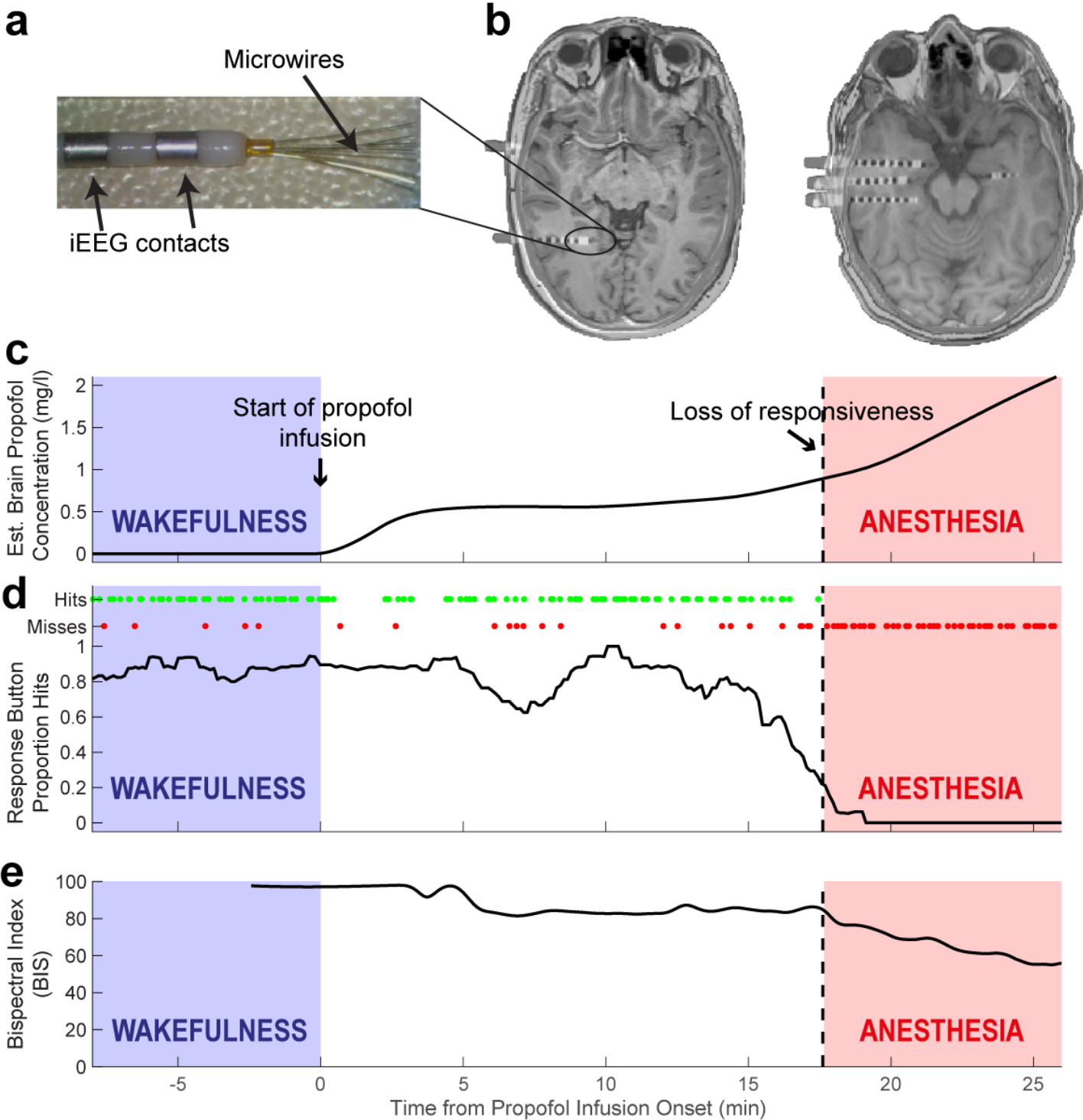
Experimental setup. (a) Depth electrodes implanted in epilepsy patients for clinical monitoring, consisting of eight 1.5mm intracranial EEG (iEEG) contacts along the shaft, and eight 40μm microwires protruding from distal tip. (b) Representative pre-implant MRI co-registered with post-implant CT used to localize electrode positions in two slices from a single subject. (c) Representative time-course of estimated propofol concentration in brain tissue (y-axis) gradually increasing during experiment. Shaded regions mark intervals of wakefulness (blue) and anesthesia (red) used for subsequent data analysis. (d) Time-course of behavioral performance on the auditory task (individual task hits/misses as green/red dots, plotted above averaged proportion of trials with hits). Vertical dotted line, loss of responsiveness. (e) Bispectral index (BIS).

We operationalized LOC as loss of behavioral responsiveness (LOR) on our task. Of note, the American Society of Anesthesiology (ASA) clinically defines such light anesthesia (characterized by unresponsiveness to verbal commands/stimuli) as “deep sedation”. While the precise propofol concentration inevitably varied between sessions, the clinical criteria for defining LOR were identical throughout, allowing us to identify neural correlates reliably associated with LOR. In each session, we compared auditory responses recorded in the first few minutes after LOR (‘anesthesia’, average bispectral index(42) BIS = 67, n = 9) with those recorded in an equally long interval during propofol-free wakefulness (‘wakefulness’, Fig. 1e, average BIS = 95).

### Anesthesia-induced LOC disrupts responses in association cortex

In wakefulness, 40Hz click-trains (Fig. 2a top & Methods) strongly entrained field potentials such that time-locked iEEG responses were highly coherent across single trials (Fig. 2a). We quantified the response fidelity of each iEEG electrode (n = 612) by calculating its Inter Trial Phase Coherence (ITPC, Methods, and Supplementary Fig. 3) at 40Hz. This measures the consistency of responses across trials on a scale of zero (random) to one (perfectly consistent). During wakefulness (Fig. 2b), significant responses (p<0.01) were found in 66% of contacts (median ITPC of all contacts = 0.40, median p = 4×10^−5^, n = 612). Responses were observed across all cortical lobes yet exhibited spatial selectivity, with temporal contacts around auditory cortex showing the strongest responses. During anesthesia, iEEG responses in association cortices were markedly reduced (Fig. 2c,d) with median ITPC dropping from 0.4 to 0.17 representing a 58% reduction. Accordingly, 65% of responding electrodes (n = 421) showed significant reduction in ITPC (p<0.05, Monte Carlo permutation test) compared to only 1.0% showing significant increase in ITPC. Robust attenuation was also evident when quantifying the distribution of gain factors between states (Methods), which was strongly skewed towards attenuation upon LOC (Fig. 2e).

**Fig. 2.**
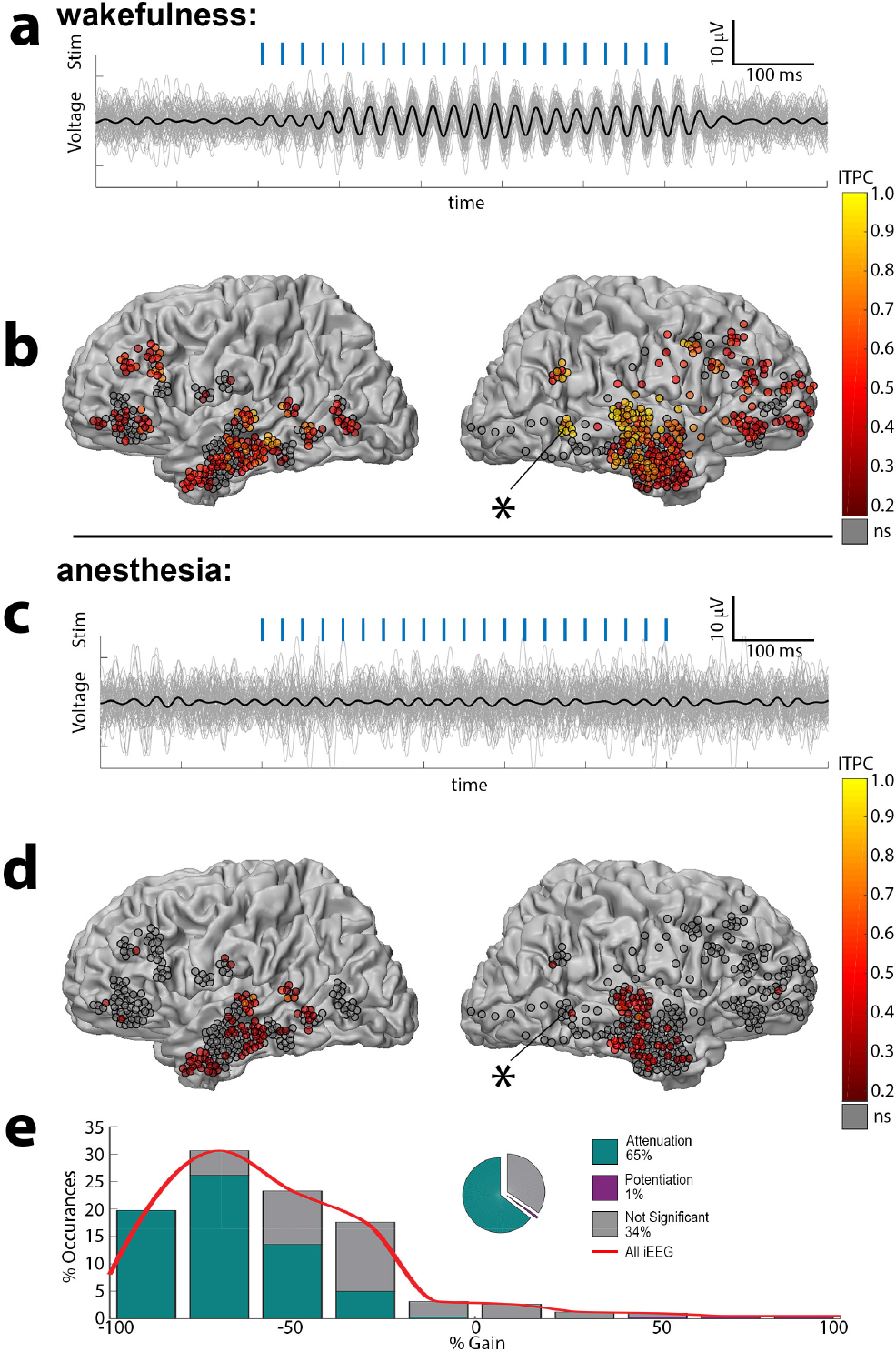
Anesthesia-induced LOC disrupts auditory responses in association cortices. (a) Responses to 40Hz click-train (stimulus, in blue) in a representative iEEG electrode in the Superior Temporal Sulcus (asterisk) during wakefulness. Thick black trace marks the average response, while thin gray traces show individual trials (b) Inter Trial Phase Coherence at 40Hz (ITPC) for each iEEG electrode (n = 612 in 9 sessions) during wakefulness, as shown on a grey-white matter boundary surface as seen from lateral view. (c) Responses of the same iEEG electrode shown in (a) during anesthesia reveal disruption of reliable responses upon LOC. (d) ITPC of the same iEEG electrodes shown in (b) revealing disruption of responses outside the primary auditory cortex. (e) Quantification of the 424 iEEG contacts showing significant responses above baseline, reveal that 65% undergo significant attenuation under anesthesia, compared to only 1% showing potentiation. The red curve represents the smoothed histogram and is used for comparison in later figures.

During anesthesia, significant (p<0.01) responses persisted in only 22% of contacts (median ITPC of all contacts = 0.17, median p = 0.20, n = 612) and 95% of these responses were observed in electrodes close to auditory cortex. Even in iEEG contacts located closest to PAC (on the shafts targeted towards the PAC), response fidelity was markedly reduced upon anesthetic-LOC (median ITPC dropped from 0.83 to 0.37, n = 27; 85% of such responsive electrodes significantly reduced ITPC compared to none showing significant increase). Disruption of iEEG responses in association cortices upon LOC was highly reproducible across individual sessions (Supplementary Fig. 4), and evident also when quantifying responses through event-related spectral power (ERSP, “induced power”, Supplementary Fig. 5).

### Robust responses persist after LOC in primary auditory cortex

In five sessions, microwires targeted seven regions located more medially in and around Heschl’s gyrus and recorded locally referenced LFPs (n = 56 microwires). Four of the seven regions (Fig. 3, regions A-D) were located in Heschl’s gyrus and exhibited short-latency (<30ms) responses (Supplementary Fig. 6). For brevity, we refer to these as “Primary Auditory Cortex (PAC)” throughout, despite the lack of systematic tonotopic mapping for delineating A1 (Methods). In contrast to disrupted iEEG responses in association cortices, PAC LFPs continued to show highly significant responses during anesthetic-LOC (Fig. 3a,b; median ITPC = 0.73, n = 32, p<10^−15^). Accordingly, the distribution of gain factors between states was largely symmetrical and only showed modest attenuation (Fig. 3c), a pattern significantly different from that found in association cortices (p<10^−12^, n = 421 in association cortex, n = 32 in PAC). Importantly, the distinction between persistent responses in PAC (median gain = −0.15, n = 32) and significant attenuation in adjacent iEEG electrodes (median gain = −0.47, n = 8) was evident when comparing these profiles in simultaneously recorded data (p = 6×10^−3^ by Wilcoxon rank-sum test, n_PAC_ = 32, n_iEEG_ = 8). In line with these ITPC results, LFP power changes in PAC microwires quantified via ERSP (Supplementary Fig. 7) showed far less attenuation than those seen in iEEG contacts across the brain (Supplementary Fig. 5, p < 10^−8^, n_PAC_ = 32, n_AsC_ = 190).

**Fig. 3.**
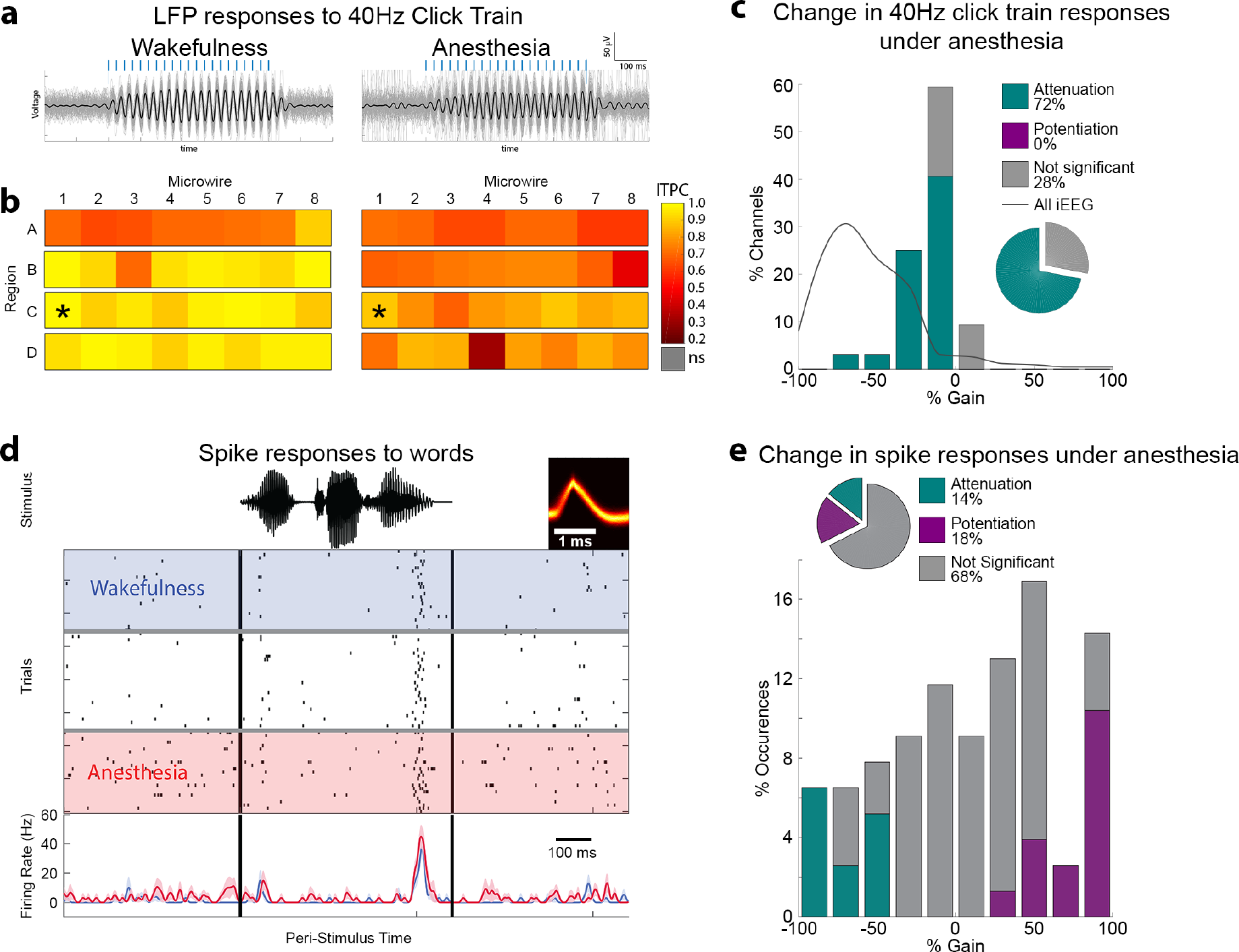
Significant responses in primary auditory cortex persist upon anesthesia-induced LOC. (a) Bandpassed responses to 40Hz click-train in a representative PAC electrode during wakefulness (left) and anesthesia (right). Thick black traces marks the average responses, while thin gray traces show individual trials. (b) Responses (ITPC) to 40Hz click-trains across all PAC microwires (n = 32, regions A-D) during wakefulness (left) and anesthesia (right) reveals significant responses persisting upon anesthesia-induced LOC. Asterisks mark the specific microwire whose responses are shown above. (c) Histogram of gain in response under anesthesia for all 32 PAC microwires shows only minimal inhibition (bars), especially when compared to the marked inhibition seen across the brain in iEEG contacts (grey curve). (d) Raster plot and peri-stimulus time histogram (PSTH, below) of representative PAC neuronal response to a word (stimulus waveform on top in black; neuron spike waveform in yellow). Rows (top to bottom) mark individual trials during deepening propofol anesthesia. Blue shading and time-course, wakefulness; Red shading and time-course, anesthesia. (e) Quantitative analysis across all PAC responses (n = 77 in 20 neurons) reveals that 68% of responses were not significantly different between wakefulness and anesthesia, 14% were attenuated in anesthesia, and 18% were potentiated in anesthesia.

Preserved PAC responses were also observed in the neuronal spiking responses to words (n = 77 responses in 20 units, see Supplementary Table 1 for details). Fig. 3d shows a representative PAC neuron, where firing rate responses were essentially unchanged between wakefulness and anesthetic-LOC (see Supplementary Fig. 8 for additional examples). A quantitative analysis across all responses (Fig. 3e, Methods) confirmed that most (68%) PAC responses did not significantly change between wakefulness and anesthetic-LOC (p>0.05, Wilcoxon rank-sum test). Similarly, components showing attenuation (14%) or potentiation (18%) were roughly equally prevalent, indicating a preservation of PAC spike responses. As was the case for relatively preserved LFP responses in PAC, spike responses in PAC significantly (p<10^−19^, n=77, n_AsC_ = 421) differed from the attenuation profile observed in association cortex iEEG responses.

### Higher-order auditory regions exhibit variable effects upon anesthetic-LOC with spike responses largely attenuated

We refer to temporal lobe regions outside PAC with higher latencies and/or weaker responses during wakefulness as higher-order auditory regions (Regions E-G, Supplementary Fig. 6, and Methods).

Region E (response latency 52ms vs. 17ms in PAC, Supplementary Fig. 6), exhibited highly reproducible auditory responses in wakefulness (Fig. 4a & 4b; median ITPC = 0.84, median p < 10^−21^, n = 8), which dramatically attenuated to noise levels after anesthetic-LOC (Fig. 4b & 4c/top; median ITPC = 0.15, median p = 0.20), as observed in association cortices (Fig. 4c/top). In line with LFP attenuation, neuronal spiking responses to words in this region (n = 56 responses in 9 units) strongly dropped around LOC. Fig. 4d shows a representative neuronal response, demonstrating clear attenuation in firing response to a word stimulus upon anesthetic-LOC (see Supplementary Fig. 8 for additional examples). A quantitative analysis across all responses (Fig. 4e) showed strong response suppression, with 84% of responses significantly attenuating (p<0.05), 16% showing no significant change, and none showing potentiation – a significantly different (p<10^−10^) profile than the preservation of PAC unit responses.

**Fig. 4.**
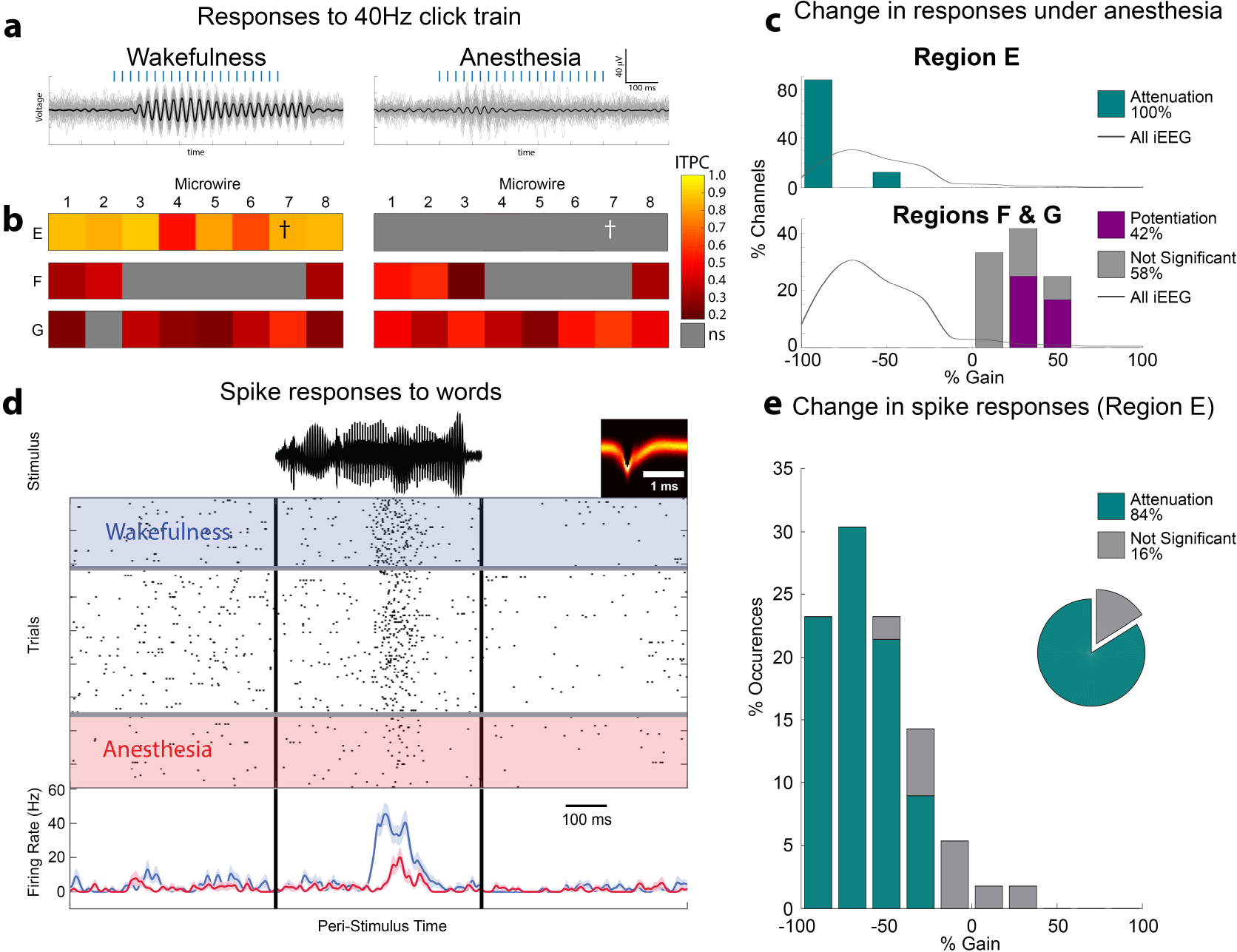
High-order auditory responses upon anesthetic-LOC. (a) Bandpass filtered responses to 40Hz click-train in a representative high-level auditory electrode during wakefulness (left) and anesthesia (right). Thick black traces marks the average responses, while thin gray traces show individual trials. (b) Responses (ITPC) to 40Hz click-trains across all high-level auditory microwires (n = 24, regions E,F,G) during wakefulness (left) and anesthesia (right). Cross marks the specific microwire whose responses are shown above, and from which the neuronal unit below (d) was recorded. Note attenuation to noise levels upon anesthesia-induced LOC in Region E, and weak potentiation of responses in Regions F & G. (c) Histograms of gain in response under anesthesia for all 20 responding higher auditory microwires (Region E, top; and Regions F & G) showing variously mainly potentiation (purple) or attenuation (cyan bars) depending on the region. This is in contrast to the marked inhibition seen across the brain in iEEG contacts (grey curve). (d) Raster plot and peri-stimulus time histogram (PSTH, below) of representative higher auditory cortex (Region E) neuronal response to a word (stimulus waveform on top in black). Rows (top to bottom) mark individual trials during deepening propofol anesthesia. Blue shading and time-course, wakefulness; Red shading and time-course, anesthesia. (e) Quantitative analysis across all the high-level auditory responses from the same Region E (n = 56 in eight neurons) reveals that 84% of responses were significantly attenuated and none were significantly potentiated. Note that this attenuation is in line with the changes in ITPC to 40Hz from the same region shown above.

Regions F & G showed a different response profile consisting of weak, albeit significant auditory responses in wakefulness (Fig. 4b; median ITPC = 0.25, n = 16, p = 3×10^−3^), which moderately strengthened upon anesthetic-LOC (Fig. 4b & 4c/bottom, median ITPC = 0.36, median p < 10^−5^). Here, 42% of microwires showed significant potentiation (p<0.05), and none showed attenuation. Anecdotally, the responses of one auditory-responsive neuron identified in these regions (Supplementary Fig. 8 bottom right) were moderately potentiated, agreeing with the LFP results. Altogether, the effects of anesthetic-LOC in higher-order auditory regions were variable (representing either inter-regional or inter-subject variability), but the strongest high-fidelity responses and corresponding spiking activities were significantly attenuated.

### Induced power responses to words during wakefulness and upon anesthetic-LOC

Induced gamma-band (40-200Hz) power is an established and widely-used proxy for neuronal activity in local neuronal populations(43). We therefore sought to measure how anesthetic-LOC affected gamma power induced by words in both iEEG and LFP signals. We identified 50 iEEG contacts and 34 microwires that showed a significant gamma response in either wakefulness or anesthesia (p<0.005, Wilcoxon signed-rank, Methods), all of which were located in the temporal lobe (Fig. 5). Figure 5a shows a representative induced power response in an iEEG electrode. Upon anesthetic-LOC, the gamma response increased its power, decreased its peak spectral frequency, and the concomitant decrease in power below 20Hz (often referred to as alpha/beta desynchronization(44)) was largely absent. A quantitative analysis across the entire dataset (Methods) confirmed that increased gamma power was a robust phenomenon across auditory-responsive electrodes (Fig. 5b,c,e; 83% increase, p < 10^−8^, n = 50 for iEEGs; 32% increase, p = 2×10^−4^, n = 34 for LFPs). Similarly, a decrease in peak gamma spectral frequency was evident across electrodes (Fig. 5b,d; 50Hz shift, p < 10^−7^ for iEEGs; 48Hz shift, p = 3×10^−5^ for LFPs). Since gamma power increase was consistently observed upon anesthetic-LOC, independently of changes in neuronal firing rates (that often showed attenuation), we examined to what extent gamma modulations caused by anesthetic-LOC were correlated with modulations in simultaneously recorded spiking responses on the same microwires. There was little or no correlation between anesthetic-LOC modulated gamma power and spiking responses (r = −0.06, n = 81). By contrast, time-locked LFP responses to 40Hz click trains as captured by ITPC were correlated with spiking responses on the same microwires (r = 0.61, n = 124). Finally, upon LOC the 10-20Hz power decrease (alpha/beta desynchronization) was consistently reduced in iEEG data (Fig. 5b, left, p = 8×10^−4^, n = 50), but a significant reduction was not observed in LFPs (Fig. 5b, right, p = 0.08, n = 34). Taken together, anesthetic-LOC causes complex changes in induced power including gamma power increase and slowing of spectral frequency profiles, which can be decoupled from local spiking activities after LOC (see also Discussion).

**Fig. 5.**
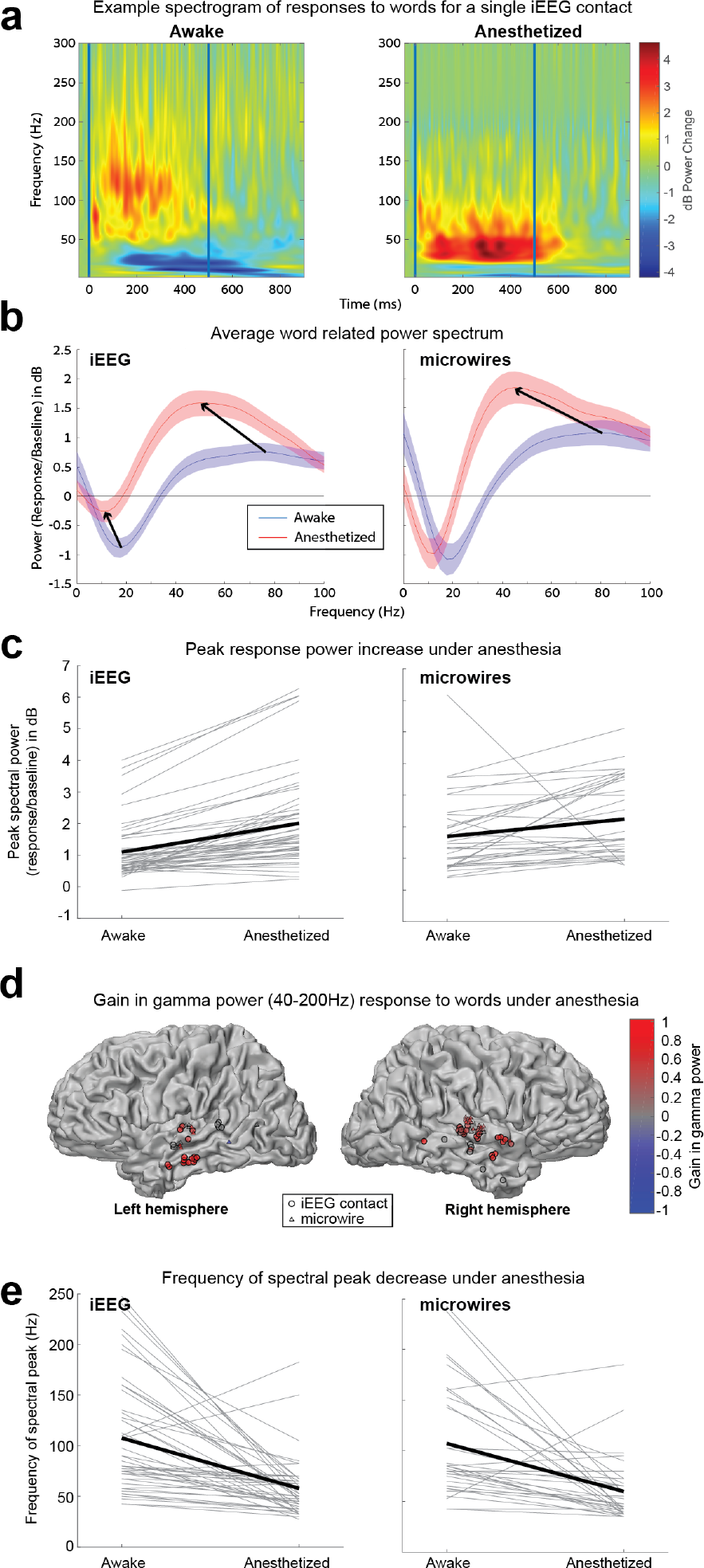
Induced gamma power responses to words upon anesthetic-LOC. (a) Example of a single temporal lobe iEEG contact event related spectral power (ERSP) averaged across words, demonstrating an increase in peak power, a reduction in the frequency of the power peak, and reduction in alpha/beta desynchronization, under anesthesia. (b) The power spectrum averaged over all 50 significantly responding (p<0.005) iEEG contacts (left) and all 34 significantly responding microwires (right), shows the same increase in peak power and shift to lower frequencies of the power peak. The iEEG spectrum also shows the reduction in alpha/beta desynchronization noted above. The peak power (c) and frequency of the power peak (d) plotted for each individual responding iEEG contact/microwire shows that these changes are consistent across contacts. (e) Plot of all 50 responding iEEG contacts and 34 responding microwires colored according to gain in gamma power under anesthesia. We see that responses are clustered exclusively around the temporal lobe, and in 83/84 case show increased power or no change.

## Discussion

By studying human auditory responses during the descent to anesthesia, we show that anesthesia-induced LOC is associated with robust disruption of auditory responses in association cortex, while significant early PAC responses persist. LOC was consistently associated with response breakdown across wide cortical territories (Fig. 2 & Supplementary Fig. 4,5). By contrast, both LFP and spike responses in PAC (Fig. 3) consistently showed robust responses after LOC. The distinction between PAC and association cortices was evident in response to both simple 40Hz click-trains and complex word stimuli. In intermediate high-order auditory regions effects were variable, and spiking responses largely attenuated around LOC (Fig. 4). Interestingly, gamma power induced by words increased after LOC while its frequency profile slowed, but such power increase did not correlate with changes in neuronal firing on the same microwires.

Since the study was conducted in neurosurgical epilepsy patients, it inherently entails some limitations that should be explicitly acknowledged. We cannot completely rule out the contribution of epileptogenic activity. In addition, the number of participants implanted with electrodes in auditory regions that volunteer to participate in research during brain surgery is limited, as is the number of recorded neurons. However, the highly consistent results across patients with different clinical profiles largely overcome these limitations. Another possible concern is that the precise brain regions targeted and the anesthesia protocols vary between sessions, as they are guided by clinical criteria. We addressed this by employing a within-session design and defining the anesthesia interval according to the same clinical endpoint (LOR) across all sessions. Furthermore, the same anesthetic agent (propofol) was used for all sessions. Future studies will determine the extent to which these findings generalize to other anesthetic agents.

In contrast to the “thalamic gating” hypothesis whereby the relay to primary sensory regions is disrupted(38), our results support the notion of preserved PAC responses under light anesthesia(29, 33, 45), as was also found for LFP responses to visual stimulation(21). Two recent studies examining the effects of propofol on human LFP auditory responses reported preserved early evoked potentials in the most posteromedial part of Heschl’s gyrus when compared to intermediate and anterolateral regions showing robust attenuation(46), as well as preservation of novelty responses over short (local deviant) time scales in auditory cortex(47). Our study extends these observations along several dimensions, most importantly by recording spike responses to complex word stimuli. The contribution of these data is two-fold: First, it consistently showed similar effects as LFP ITPC results, thereby allowing to extrapolate and confidently interpret time-locked LFP responses as reflecting local spiking activities even when these were not available. Second, it exposed interesting differences after LOC compared with simultaneously recorded gamma power changes. The fact that spiking responses are preserved in PAC is of particular importance since this demonstrates unequivocally that during anesthetic-LOC not only is the synaptic input into PAC preserved, but also the circuit’s output. The apparent discrepancy with studies reporting attenuation of PAC responses under anesthesia(26–28) may reflect effects of deep surgical anesthesia unrelated to LOC, and highlights the importance of using “just-hypnotic” anesthesia when studying LOC. While the present results establish preserved PAC responses to isolated stimuli upon anesthesia-induced LOC, they do not preclude potential effects on select neuronal populations, precise spike timing, or contextual effects that require integration across long temporal intervals.

Attenuated EEG responses to 40Hz click-trains have been repeatedly demonstrated upon anesthesia-induced LOC and have been suggested as a useful marker for assessing the depth of anesthesia(31, 48). Our results indicate that disruption of scalp EEG responses represent attenuated activity in association cortices, as can be expected given that preserved PAC responses only modestly contribute to total cerebral activity, and given the medial location of PAC that is less accessible to non-invasive scalp EEG. Indeed, our own iEEG results show robust attenuation upon anesthesia-induced LOC (Fig. 2). The effectiveness of the specific 40Hz click-train stimulus in revealing correlations with LOC may be related to an inability of large neuronal populations to reliably synchronize coherently with high-frequency stimulation in disconnected states(49). Studies in natural sleep demonstrate a similar distinction between preserved tracking of acoustic properties and PAC responses(50) vs. high-order activity in association cortex(51). Thus, a functional disconnection between primary sensory and association cortex may be a general property of LOC not only due to anesthesia, as demonstrated also by direct perturbation of cortical activity with transcranial magnetic stimulation (TMS)(52–54). In addition, our results join findings from other lines of research showing that when sensory stimuli are not perceived, activity in primary sensory regions is largely preserved whereas activity in association cortices is attenuated(55–58).

Our findings reveal complex effects on induced power changes, where anesthetic-LOC was associated with gamma power increase and a slowing of frequency profiles (Fig. 5). Such changes are in line with previous studies showing that anesthesia/sedation increases gamma responses power in-vitro(59) and in humans in-vivo(60, 61). Similarly, a slowing of spectral frequencies upon anesthesia has been observed in-vitro(59, 62), and in monkeys(63) and humans in-vivo(61). Whilst induced gamma power usually accompanies neuronal spiking activity(43), our results show that the relation between induced gamma power and local spiking activity may change across states. The extent of potential decoupling between gamma activity and neuronal firing remains unclear, and we can only speculate about its underlying source. One possibility is that these two measures emphasize the contribution of different neuronal populations that diverge upon anesthetic-LOC. For example, gamma oscillations are closely coupled with inhibitory GABAergic interneurons(64), whilst extracellular recordings mostly capture excitatory pyramidal cells(65). Differences may also stem from differential contributions from distinct cortical layers or cell populations at different spatial scales. We also find that stimulus-induced decrease in 10-20Hz power (prevalent during wakefulness) diminishes upon anesthetic-LOC (Fig. 5). Given that alpha/beta desynchronization has been linked with top-down signaling (66), this may support the notion that anesthesia predominantly affects top-down signaling (29, 67).

What could be the mechanisms by which LOC disrupts cortical functional connectivity? As noted in the introduction, anesthesia may act at multiple subcortical mechanisms simultaneously(2, 9, 11–13, 15, 68) in a manner that ultimately results in reduced cortical connectivity. For example, anesthesia may exert its effects by affecting endogenous subcortical sleep/wake neuromodulation centers that modify thalamocortical activity(2, 13). Accordingly, the distribution of modulatory receptors may differ between primary sensory regions and association cortices, rendering the latter unresponsive to preserved ascending input. An alternative explanation for the inability to effectively drive responses in association cortices may reflect a disruption to indirect cortico-thalamo-cortical pathways linking cortical regions via high-order thalamic nuclei(69, 70). Thus, refining the traditional “thalamic gating” notion from disruption of primary relay nuclei, to focus on high-order thalamic nuclei, could reconcile the present results with the well-established effect of anesthesia reducing thalamic activity(38). Alternatively, our findings may reflect a “multiple hit mechanism”, whereby anesthesia causes multiple small perturbations at each step in the sensory processing chain. The sum of these perturbations may seem modest in the PAC as measured with current methods, but may have accumulated sufficiently to be noticeable in the higher association cortices. Finally, anesthetics could affect top-down signaling that may be more dominant in association cortices(67, 71), possibly involving specific ion channels that link bottom-up and top-down signaling(72). Furthermore, anesthetic agents may employ several of these mechanisms in parallel since they reach much of the brain and central nervous system almost simultaneously. Beyond the specific mechanisms at play, the disruption of effective cortical connectivity is consistent with a number of theoretical perspectives(73, 74).

In conclusion, our findings demonstrate that anesthesia-induced LOC disrupts auditory responses beyond primary cortex while significant responses persist in PAC. The fact that the major disruption in sensory signaling upon LOC occurs after primary cortices highlights impaired effective cortical connectivity as a key factor in LOC, a factor that should guide future mechanistic research of LOC, development of anesthetic agents, and design of depths of anesthesia monitoring schemes.

## Materials and Methods

### Subjects

Eight patients (six male, see Supplementary Table 1 for details) with intractable drug-resistant epilepsy were implanted with chronic depth electrodes, or subdural cortical grids and strips (one session), for 7-10 days, to identify seizure foci for potential surgical treatment. Electrode location and anesthesia management (below) were based solely on clinical criteria. One patient was implanted on two different occasions due to inconclusive clinical results from the first hospital admission. All patients provided written informed consent to participate in the research study, under the approval of the Institutional Review Board at Tel Aviv Sourasky Medical Center, or the University of Bonn Epileptology Department (one individual). Data were collected prior to removal of the electrodes under anesthesia at the end of monitoring on the medical ward.

### Anesthesia

Standard anesthesia monitoring was performed according to the guidelines set by the American Society of Anesthesiologists (ASA), complemented with bispectral index monitoring(42) (BIS VISTA™ Monitoring System, Medtronic, Boulder, CO, USA). Propofol was infused using a simple (non-Target Controlled Infusion) syringe pump, with infusion rates increasing in a step-wise fashion over 20-40 minutes (to a maximum of 45 to 400 μg/kg/min depending on the patient and according to the judgement of the anesthesiologist) to achieve anesthesia whilst maintaining spontaneous ventilation. In two sessions (see Supplementary Table 1) adjuvant agents (fentanyl, midazolam or remifentanil) were given according to clinical considerations. When a sufficient depth of anesthesia was achieved (aiming to avoid movement in response to painful stimulus whilst maintaining spontaneous ventilation), the neurosurgeons removed the electrodes. Post-hoc pharmacokinetic and pharmacodynamic modelling using the three-compartment Marsh Model(75–78) (Supplementary Fig. 1), were used to estimate the momentary effect-site concentration of propofol in brain tissue (Fig. 1 and Supplementary Fig. 2).

### Auditory Paradigm

Auditory stimuli were continuously presented throughout the descent to anesthesia (20 – 70 minutes) using headphones (8 sessions) or speakers (1 session), with sound intensity levels adjusted at the start of each session to be comfortably audible. 40Hz click-trains (duration = 500ms, 21 clicks) were presented in all recording sessions. Additionally, up to seven word stimuli (duration = 500ms - 600ms), including one task target (duration 500ms), were presented in seven sessions. Auditory stimuli were presented in a pseudorandom order, at 1.5s intervals (± 400ms jitter). Patients were instructed to press a response button upon hearing a specific target word, to determine the moment of loss of responsiveness under anesthesia.

### Electrophysiology

In eight sessions, patients had been implanted with 5 – 11 depth electrodes targeting different brain regions (Fig. 1B), whereby each electrode consisted of eight platinum iEEG contacts along their shaft, eight additional microwires protruding a few mm from the distal tip, and a 9^th^ low-impedance reference microwire(79). In one of these sessions only iEEG data was available. In a separate ninth session, a patient was implanted with a combination of two two-dimensional subdural electrocorticography (ECoG) grid arrays and four one-dimensional subdural ECoG strips (i.e. instead of depth electrodes). Data were recorded using either Blackrock (Salt Lake City, UT, USA) or Neuralynx (Bozeman, MT, USA) data acquisition systems. Microwire data were sampled at 30kHz (Blackrock) or 32,768 Hz (Neuralynx) and referenced to the local low impedance 9^th^ microwire on each electrode, or in one session to a single low impedance 9^th^ microwire (session 6, see Supplementary Table 1). iEEG and ECoG data were sampled at 2 kHz (Blackrock) or 2,048 Hz (Neuralynx) and referenced to a central scalp electrode (Cz).

### Electrode Localization

Pre-implant MRI scans (Siemens Prisma scanner or GE Signa scanner, 3 Tesla, T1 sequence, resolution 1mm × 1mm × 1mm) were co-registered with post-implant CT scans (Philips MX8000 or Brilliance, resolution 1.5mm × 0.5mm × 0.5mm or 0.75mm × 0.5 mm × 0.5mm) to identify locations of electrodes. Individual subject data were further transformed into Talairach space(80) to facilitate simultaneous visualization of electrode positions in different individuals. To this end, electrode locations were marked with 5mm spheres on the standard brain, and intersections with the grey-white matter boundary surface were identified. The cortical surface was then inflated and flattened by unfolding and cutting along the calcarine sulcus and predefined medial anatomical landmarks (Supplementary Fig. 4). When plotted on folded grey-white matter boundary surface (Fig. 2, Fig. 5 & Supplementary Fig. 5), the location of each electrode was defined as the center of mass of the intersection of the 5mm sphere with the surface, and a circle was drawn in that location (minimal jitter was introduced to allow visualization and avoid hidden data).

### Defining “Wakefulness” vs. “Anesthesia” Periods

To compare auditory responses during propofol-free wakefulness to those under light anesthesia (formally defined by the ASA as “deep sedation”: unresponsiveness to simple verbal instructions) we defined the anesthesia period as the first eight minutes following LOR. In order to avoid excessively deep anesthesia, we confirmed that BIS values remained above 50(42). Supplementary Table 1 and Supplementary Fig. 2 provide full details regarding recording durations and definitions of wakefulness and anesthesia intervals in each session. LOR was defined as the midpoint between the last correct behavioral response and the following missed target word. Consideration was given to assessing depth of anesthesia according to responses to intermittently delivered, graded severity of stimuli (e.g. the Observer’s Assessment of Alertness/Sedation Scale OAA/S (81) which involves “mild prodding/shaking”), however this itself can wake subjects from shallow states of unconsciousness (82), and give a false measurement of alertness, and so we chose to rely on unprompted performance in the word identification task described above. In Sessions 1, 2, and 7 where it was not possible to define LOR based on behavioral button press data, we defined LOR as the time when the patient was observed to stop reacting purposefully to his environment, by noting for example when he stopped talking to care-givers, or when he responded only to painful stimuli such as scalp scrubbing or needle sticks. For each experimental session, we chose an interval of propofol-free wakefulness with identical duration for a within-session comparison. In two sessions the anesthesia period was truncated to avoid deep anesthesia (sessions 7 & 8), defined by BIS < 50. Whenever eight minutes were not available for either wakefulness or anesthesia, we used the longest possible noise-free intervals with identical durations in both wakefulness and anesthesia (Supplementary Fig. 2 & Supplementary Table 1). Average BIS values quoted in the text were calculated by integrating over BIS curves during wakefulness or anesthesia for each session (with linear interpolation between known data points), and then taking the median across sessions. In all but one session, patients breathed spontaneously and maintained their airways throughout the period defined as anesthesia. In Session 2 the patient required a jaw thrust and Guedel oral airway to maintain his airway.

### Analysis of iEEG/LFP responses to 40Hz click trains

Responses to 40Hz click trains were quantified using Inter-Trial Phase Coherence (ITPC, Fig. 2 & 3), also known as the “phase-locking factor”(83, 84):

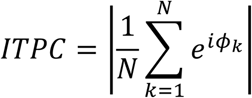

where N signifies the number of trials, and *ϕ*_k_ the phase of the spectral estimate for trial k for the 40Hz frequency.

The statistical significance values associated with ITPC were estimated as in Zar (1999)(85) with a significance threshold of α = 0.01:

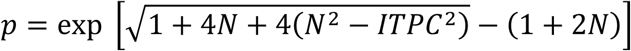

where N is the number of trials and ITPC is the calculated ITPC.

Quoted averages of ITPC p-values are the median of these p-value estimates. Median ITPC values for Supplementary Fig. 4 were calculated on the 421 iEEG electrodes showing significant ITPC in either wakefulness or anesthesia.

The gain factor between wakefulness and anesthesia was calculated as in as in(50, 86):

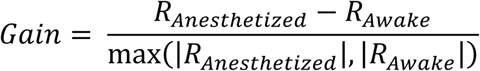

where R_Anesthetized_ and R_Awake_ is the ITPC for a particular channel during wakefulness or anesthesia. We obtained largely similar results when calculating gains using only R_Awake_ in the denominator (not shown). Additionally, responses were quantified by calculating the Event-Related Spectral Power (ERSP) at 40Hz (Supplementary Fig. 5 & 6), reflecting power modulations that are not necessarily phase-locked and known also as “induced power”. To this end, the 40Hz spectral power was calculated by applying a Hann window over the 500ms of the response (or baseline) iEEG/LFP signal, and calculating the squared 40Hz amplitude component of the Fourier transform. Spectral power during baseline was subtracted from that during response, and the result divided by the median baseline spectral power at 40Hz. This ERSP was averaged across all trials for each electrode (Supplementary Fig. 5 & 6). Statistical significance of ERSP responses was calculated using a Wilcoxon signed-rank test, with significance level of α = 0.01 (non-significant values are masked in grey in Supplementary Fig. 5).

### Response latency

Latency (Supplementary Fig. 6b) was calculated based on the locally referenced LFP response to the 40 Hz click train during wakefulness for each individual microwire targeted around Heschl’s Gyrus (Fig. 3 & 4). This method requires responses to be strongly coherent between trials, and was possible to implement in 72% of auditory-responsive electrodes with ITPC > 0.5 (most of which were > 0.85). The mean LFP response ([−600,2000]ms from stimulus onset) to 40 Hz click train was bandpassed at 30-150Hz (4^th^ order zero-phase-shift digital butterworth filter) to include the LFP response at 40 Hz and its harmonics. Then, its envelope was extracted using the Hilbert transform. Using a 525 ms length moving window (window size was chosen to match the LFP response to the stimulus of 21 clicks with 25 ms ISI), latency was defined as the onset time of the window where the largest mean envelope was obtained.

### LFP induced power response to words

Responses were quantified by measuring, for each microwire and iEEG contact, the event-related gamma power (40-200Hz, mean of FFT magnitudes squared) for each trial across words (100-500ms Hann window, post-trial onset) and comparing it with power during the corresponding pre-stimulus baseline period (−450 to −50 ms). “Responding channels” were defined as those in which this gamma power during response was significantly different from baseline gamma power (p<0.005, Wilcoxon signed-rank), and were selected for further analysis. Response power spectra for each responding channel were calculated by dividing the event-related power in each trial (100-500ms) at each frequency, by the corresponding baseline power, and then taking the median across wakefulness (or anesthesia) trials. This spectrum per channel was then averaged (mean ±SEM) across iEEG or microwire channels to visualize the grand mean profile (Fig. 5b). The peak spectral power during wakefulness or anesthesia was determined for each channel separately by identifying the highest peak in the power spectrum, and finding its power (Fig. 5c) and frequency (Fig. 5e). The alpha/beta desynchronization trough power during wakefulness or anesthesia was determined for each channel by identifying the deepest trough in the power spectrum that was to the left of (lower frequency than) the peak power above, and which also lay below 50Hz. On occasions where no such trough was identified, the alpha/beta desynchronization was assumed to have disappeared, and the trough value was set to zero dB. Gain factors between wakefulness and anesthetic-LOC for gamma power (Fig. 5d) were calculated in each channel separately as follows. Event-related gamma power change compared to baseline (as described above) was normalized by dividing by the median baseline power across trials, and averaged over awake (or anesthesia) trials, yielding a “gamma power response” for that channel during wakefulness (or anesthesia). The gain factor was then calculated as described above for ITPC, but substituting R_Anesthetized_ and R_Awake_ with the gamma power response for a particular channel during wakefulness or anesthesia.

### Analysis of neuronal spiking activity

Spike sorting was performed using the wave_clus toolbox for Matlab as described previously(87, 88): (i) extracellular microwire recordings were high-pass filtered above 300Hz, (ii) a 5 × SD-approximation (*median*{|*data*|/0.6745}) threshold above the median noise level was computed, (iii) detected events were clustered using superparamagnetic clustering, and categorized as noise, single- or multi-unit clusters. Classification of single- and multi-unit clusters was based on the consistency of action potential waveforms, and by the presence of a refractory period for single units, i.e. less than 1% of inter-spike-intervals (ISIs) within 3ms.

Raster and peri-stimulus time histograms (PSTH) were produced for each neuronal unit in response to each word (see Supplementary Fig. 8 for examples). Instantaneous firing rate traces (all PSTHs below raster plots) were calculated by smoothing the binary spike trains with a Gaussian kernel (σ = 5 ms). Raster plots were inspected manually, and periods of increased firing above baseline were identified as “response components” (black arrows in Supplementary Fig. 8). A single unit responding to a few isolated acoustic events (e.g. phonemes) within a single word stimulus could therefore have one or more response components (Supplementary Fig. 8), and the firing rate (total spikes) of each such component was compared separately between wakefulness and anesthesia. Categorization of response components as potentiated, attenuated or unchanged was determined via a Wilcoxon rank-sum test with a significance level of α = 0.05.

### Statistics

P-values were calculated using Wilcoxon rank-sum tests, Wilcoxon signed-rank tests, or Monte Carlo permutation tests, as reported above, while treating iEEG or LFP for different electrodes as independent measures. Error bars in all figures denote standard error of the mean (SEM = SD/√n, where n is the number of data points) unless otherwise stated.

## Acknowledgements

We thank the patients for their cooperation. Rivi Radomsky, Sari Nagar and Noa Regev for administrative help. Netta Neeman for help localizing electrodes. Margaret Ekstein and Thomas Reber for help with anesthesia research. Israel Nelken, Rafael Malach, Aeyal Raz, and members of Nir lab for discussions and suggestions.

## Author contributions

I.F. and Y.N. conceived of research.

A.M. and A.J.K designed experiments and analyzed data.

A.J.K, A.M., H.G.S, A.T., H.H., F.M. and Y.N. collected data.

I.F., I.S., and J.B. performed surgeries.

D.H., M.S., and I.M. supervised anesthesia.

F.F. supervised clinical care and analyzed epilepsy profiles.

A.J.K, A.M., H.G.S, I.F. and Y.N. wrote the manuscript.

All authors provided ongoing critical review of results and commented on the manuscript.

## Data and code availability

The data that support the findings of this study and the computer code used to generate the results are available from the corresponding author upon reasonable request.

