## Supplementary Material for "Anesthesia-induced loss of consciousness disrupts auditory responses beyond primary cortex"

| **Patient** | 1 | 2 | 3 | 4 | | 5 | 6 | 7 | 8 |
|---|---|---|---|---|---|---|---|---|---|
| **Session** | 1 | 2 | 3 | 4 | 5 | 6 | 7 | 8 | 9 |
| **Age** | 23 | 40 | 49 | 24 | 25 | 31 | 40 | 23 | 39 |
| **Gender** | male | male | male | female | female | female | male | male | male |
| **Weight (kg)** | 73 | 88 | 62 | 45 | 45 | 97 | 64 | 110 | 70 |
| **Recording duration (min)** | 41 | 59 | 90 | 79 | 53 | 46 | 50 | 48 | 40 |
| **Duration of Wakefulness / Anesthesia (sec)** | 332 (limited by length of noise-free wakefulness recording) | 480 | 300 (limited by length of noise-free wakefulness recording) | 465 (limited by length of noise-free anesthesia recording) | 480 | 480 | 177 (limited by rapid anesthetic descent) | 390 (limited by rapid anesthetic descent) | 480 |
| **Stimuli** | 40Hz CT | 40Hz CT | 40Hz CT; target + 3 other words | 40Hz CT; target + 6 other words | 40Hz CT; target + 5 other words | 40Hz CT; target + 4 other words | 40Hz CT; target + 4 other words | 40Hz CT; target + 4 other words | 40Hz CT; target + 4 other words |
| **Behavioral Task** | no | no | yes | yes | yes | yes | yes | Yes | yes |
| **Anesthetic Agents** | propofol; fentanyl; midazolam | propofol; remifentanil | propofol | propofol | propofol | propofol | propofol | propofol | propofol |
| **Auditory region label (Fig. 3)** | -- | A | -- | B | E | C (Lt) & D (Rt) | N/A (grid) | -- | F & G |
| **No. of auditory responsive units** | 0 | 0 | 0 | 1 SU / 2 MU | 5x SU / 3x MU | 10x SU / 7x MU | N/A (grid) | N/A (no microwire data) | 1x MU |
| **Total No. of units from wires in/around Heschl's Gyrus** | 0 | 0 | 0 | 1x SU / 8x MU | 6x SU / 3x MU | 11x SU / 14x MU | N/A (grid) | N/A (no microwire s) | 3x SU / 4x MU |
| **Regions targeted by most medial contacts** | RAH, RA, RMF, LAH, LA, LMF | RA, RAH, REC, RHSC, RPHG, RMF, LA, LAH, LHSC, LMF | LA, LAH, LMTO, LOp, LOm | RMF, REC, RMH, RA, RSTG, RAC, RpSMA, LEC-LPHG, LAH, LAC, LpSMA | RAH, RaSTG, RmSTG, RmP, RpSMA, RdAC, RmSFG, RMF, RmOF, RFP, RMTO | LAH, RAH, LA, RA, LEC, REC, LHSC, RHSC, LMH, LPHC, RPHC | RFTG, RBTOS, RPBTS, RIFS, RMFS, RFSG | LAF, LdAC, LvaCING, LSF, LpGaCIN G, LMH, LEC, RdAC, RpGaCIN G, RSF | LOF, LAI, LMI, LSI, LPI, LSTG, LPHG, RA, REC, RMH |
| **Seizure Onset Zone** | RAH | Predominant ly middle RHSC, but also middle/latera l LHSC | LAH | Right temporal / frontal lobes | Right superior frontal gyrus | Left superior temporal gyrus near LHSC | Poorly defined. Probably in Rt anterior temporal lobe | LAF/ LpGaCIN G | Seizure onset zone unclear |

### Table S1. Data acquisition details

Data acquisition details for the nine recording sessions included in this study. Rows (top to bottom) show patient number, session number, age, gender, weight, recording duration, auditory stimuli, behavioral task, anesthetic agents, BIS monitoring, label of auditory region, number of auditory responsive units, targeted regions, and seizure onset zone. Sessions 1 – 6 and 8 - 9 involved depth electrodes whilst session 7 involved a subdural grid and subdural strips. Sessions 4 and 5 involved the same patient who was re-admitted with new electrode locations due to inconclusive clinical results from her first hospital admission. Abbreviations: 40Hz CT = 40Hz click-train; Lt=left; Rt=right; SU = single neuronal unit; MU = multi neuronal unit; L=left hemisphere; R = right hemisphere; AH=anterior hippocampus; A=amygdala; MF=medial frontal; MTO = medial temporal occipital junction; Op = occipital lobe, posterior aspect; Om= occipital lobe, medial aspect; EC=entorhinal cortex; HSC=Heschl's gyrus; MH=middle hippocampus; STG=superior temporal gyrus; AC = Anterior cingulate cortex; pSMA=pre-Supplementary Motor Area; PHG = parahippocampal gyrus; aSTG = superior temporal gyrus – anterior part; mSTG = superior temporal gyrus – middle part; mP = parietal lobe – medial aspect; dAC = dorsal anterior cingulate; mSFG = middle superior frontal gyrus; mOF = medial orbitofrontal; FP = frontal pole; MC = middle cingulate gyrus; PHC = parahippocampal cortex; FTG = frontotemporal grid; BTOS = basal temporal occipital strip; PBTS = post-basal temporal strip; IFS = inferofrontal strip; MFS = middle frontal strip; FSG = frontal small grid; AF = anterior frontal; vaCING = ventral anterior cingulate; pGaCING = pre-genual anterior cingulate; SF = superior frontal

**a**

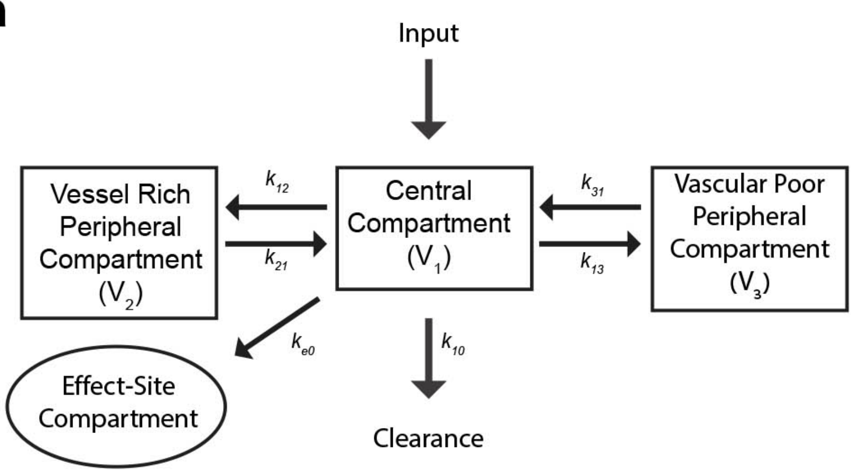


**b**

$$\frac{dC_e}{dt} = k_{e0} \times \frac{m_1}{V_1}$$
$$\frac{dm_{clearance}}{dt} = k_{10} \times m_1$$
$$\frac{dm_2}{dt} = k_{12} \times m_1 - k_{21} \times m_2$$
$$\frac{dm_3}{dt} = k_{13} \times m_1 - k_{31} \times m_3$$
$$\frac{dm_1}{dt} = input - \frac{dm_2}{dt} - \frac{dm_3}{dt} - \frac{dm_{clearance}}{dt}$$

$$V_1 = 0.228\,\mathrm{L\,kg^{-1}} \times PntWeight$$
$$V_2 = 0.463\,\mathrm{L\,kg^{-1}} \times PntWeight$$
$$V_3 = 2.893\,\mathrm{L\,kg^{-1}} \times PntWeight$$
$$k_{10} = 0.119\,\mathrm{min^{-1}}$$
$$k_{12} = 0.112\,\mathrm{min^{-1}}$$
$$k_{13} = 0.042\,\mathrm{min^{-1}}$$
$$k_{21} = 0.055\,\mathrm{min^{-1}}$$
$$k_{31} = 0.0033\,\mathrm{min^{-1}}$$
$$k_{e0} = 0.26\,\mathrm{min^{-1}}$$

**Fig. S1. Modelling time dynamics of propofol concentration**

(a) Three compartment pharmacokinetic model ( “Marsh Model” (1)) used to estimate propofol concentration in the effect-site compartment (analogue to the brain) of each patient at any given moment (Fig. 1c and Supplementary Fig. 2) given the known rate of propofol infusion (“Input”), and parameters. (b) Actual parameter values used, based on patient’s weight (PntWeight) (2–4). Note that while V2 and V3 do not explicitly appear in the model equations, they ensure that concentrations in the various compartments (= mass / volume) are equal during equilibrium. The effector site concentrations estimated by this model at the moment of LOR ranged between 0.49 – 2.8 mcg/ml (for patients who did not receive adjuvant agents). Similar (albeit slightly higher) estimates were obtained when using the Schnider model (not shown), ranging between 0.91 – 4.2 mcg/ml. Such variability in effector site concentrations at LOR is in line with literature (5, 6).

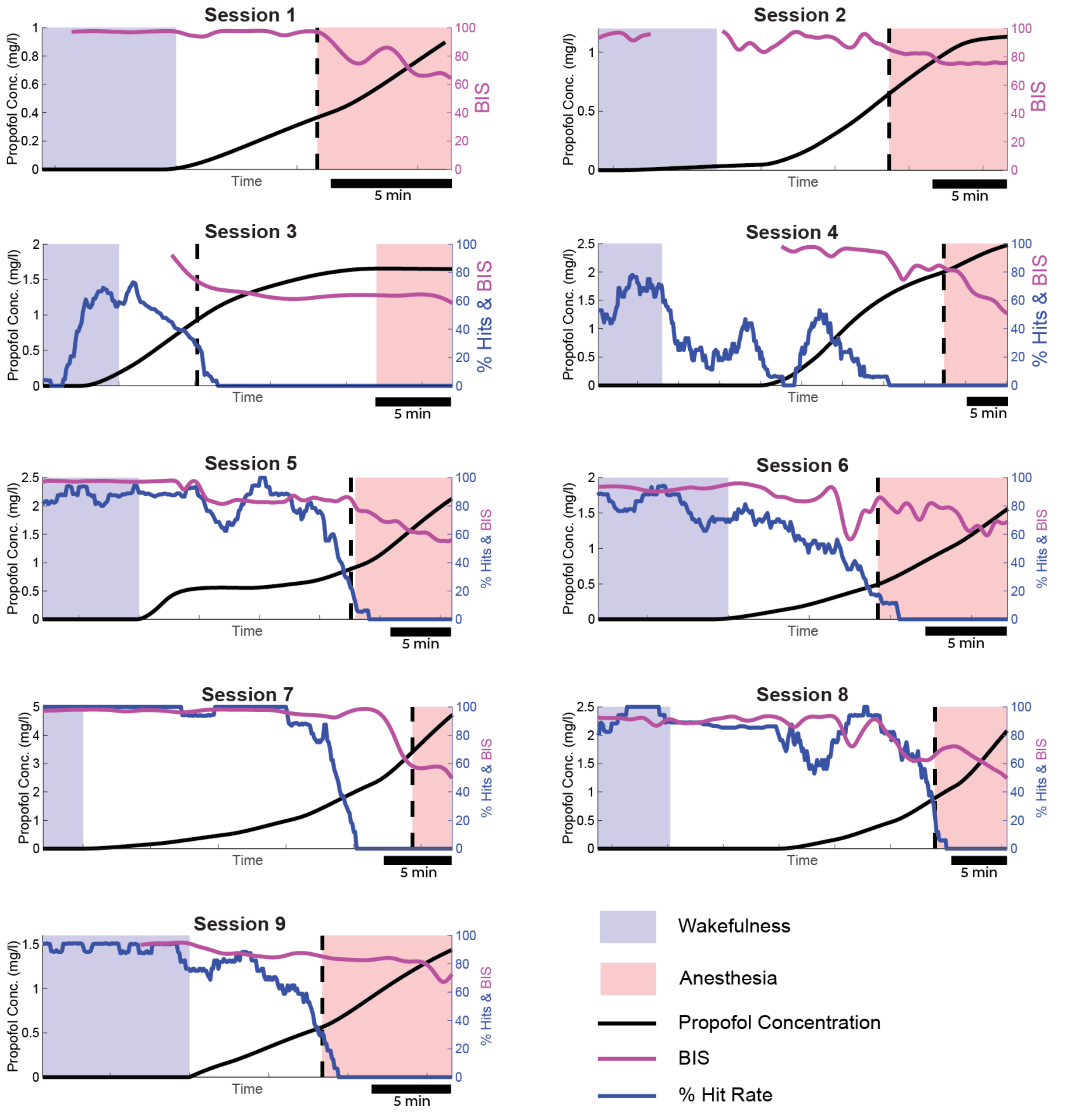


**Fig. S2. Time dynamics of anesthesia, behavior, and BIS in each experimental session**

Superimposed time dynamics in each experimental session (subpanels) showing estimated effector site (brain) propofol concentrations (black trace and y-axis on left), BIS values (purple trace and y-axis on right), and behavioral performance (blue trace and y-axis on right, proportion of successful button presses). The vertical dashed black line marks loss of responsiveness (the final time the patient successfully pressed the button to the target word). The behavioral performance ("%Hits") curve is calculated as a moving window average over approximately 16 trials (≈150 sec), and therefore may overshoot the loss of responsiveness line. Blue and red shading mark periods of wakefulness and anesthesia, respectively, used for analysis of auditory responses (whenever possible, about 8 minutes of light anesthesia with BIS>50, Methods). In session 3, the anesthesia period for analysis occurred 12 minutes after LOR due to technical issues before that time.

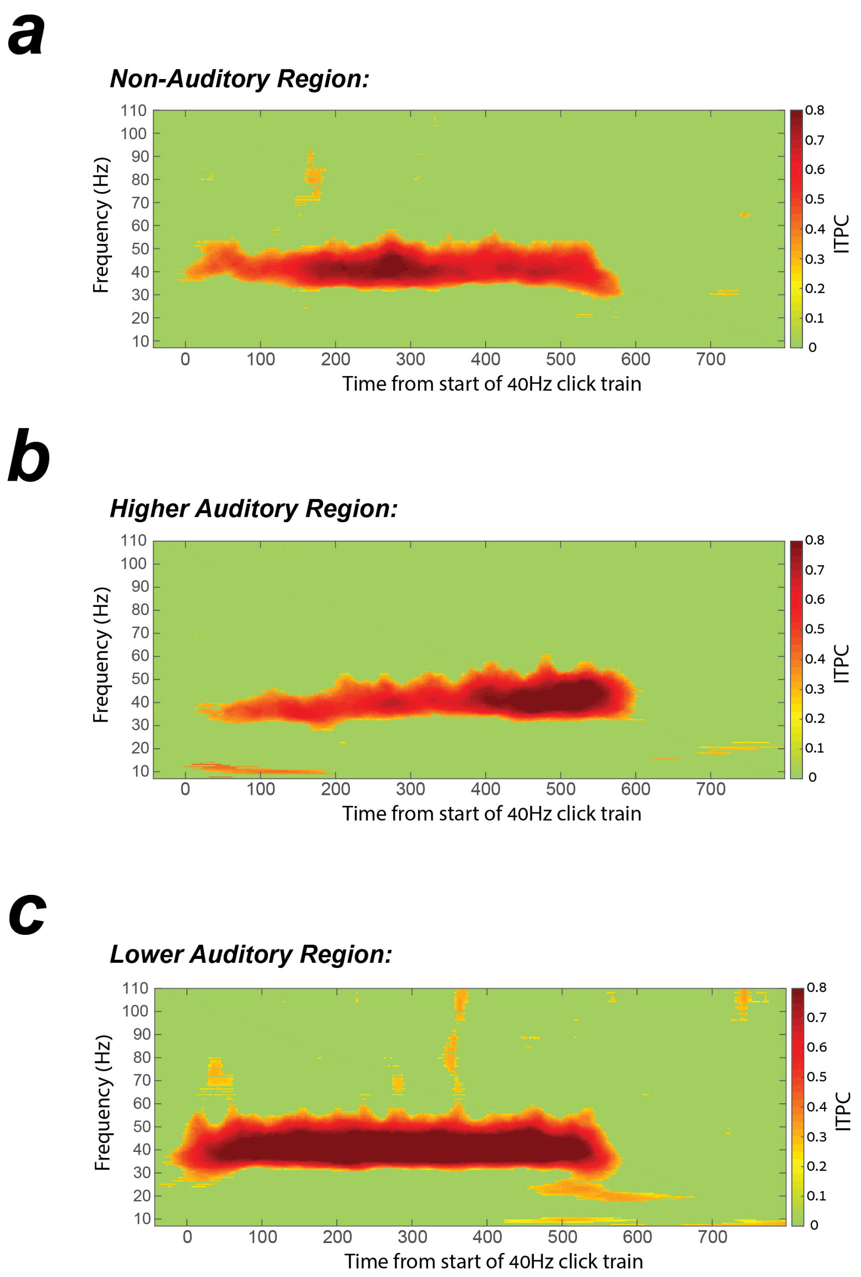


**Fig. S3: 40Hz click-trains elicit significant Inter Trial Phase Coherence (ITPC) at stimulation frequency**

Wideband ITPC spectrogram of responses to 40Hz click-trains during wakefulness in (a) a representative iEEG macro-electrode in association cortex (session 5, 64 trials), in (b) a representative LFP microwire electrode in higher auditory region (session 5, 64 trials), and in (c) a representative LFP microwire electrode in the PAC (session 6, 75 trials).

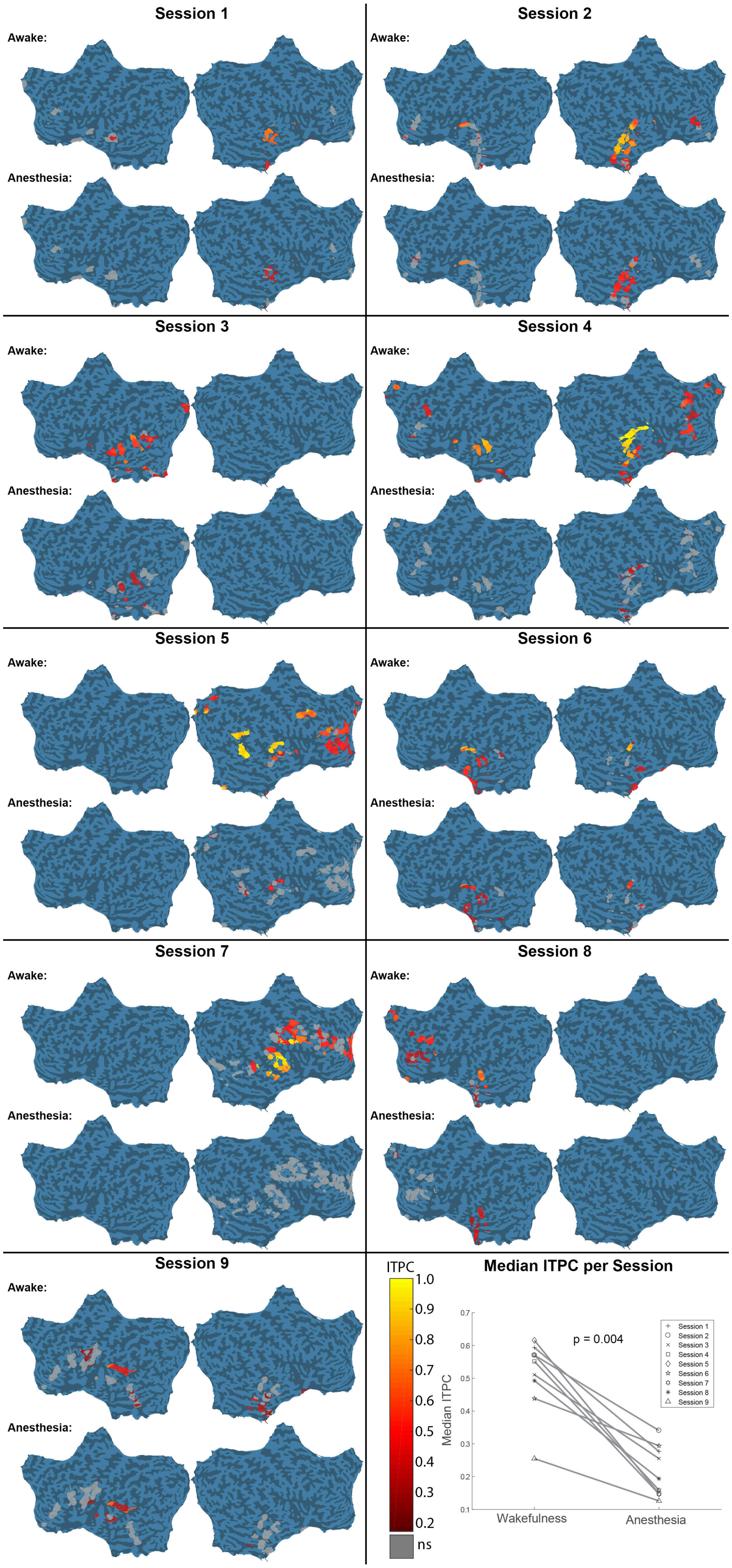
Session 1
Awake:
Anesthesia:
Session 2
Awake:
Anesthesia:
Session 3
Awake:
Anesthesia:
Session 4
Awake:
Anesthesia:
Session 5
Awake:
Anesthesia:
Session 6
Awake:
Anesthesia:
Session 7
Awake:
Anesthesia:
Session 8
Awake:
Anesthesia:
Session 9
Awake:
Anesthesia:
ITPC
1.0
0.9
0.8
0.7
0.6
0.5
0.4
0.3
0.2
ns
Median ITPC per Session
p = 0.004
Median ITPC
0.7
0.6
0.5
0.4
0.3
0.2
0.1
Wakefulness
Anesthesia
Session 1
Session 2
Session 3
Session 4
Session 5
Session 6
Session 7
Session 8
Session 9

**Fig. S4: Attenuation of iEEG responses outside primary auditory cortex in individual sessions**

Each subpanel shows auditory responses in iEEG macroelectrodes in a separate session (see Supplementary Table 1 for details). Each circular patch shows a specific iEEG electrode and its color (colorbar) denotes Inter Trial Phase Coherence at 40Hz (ITPC) during wakefulness (top) and during anesthesia (bottom), as shown on a standard flat cortical surface. Bottom-right panel shows median ITPC for all significant iEEG contacts, per session in wakefulness vs. anesthesia.

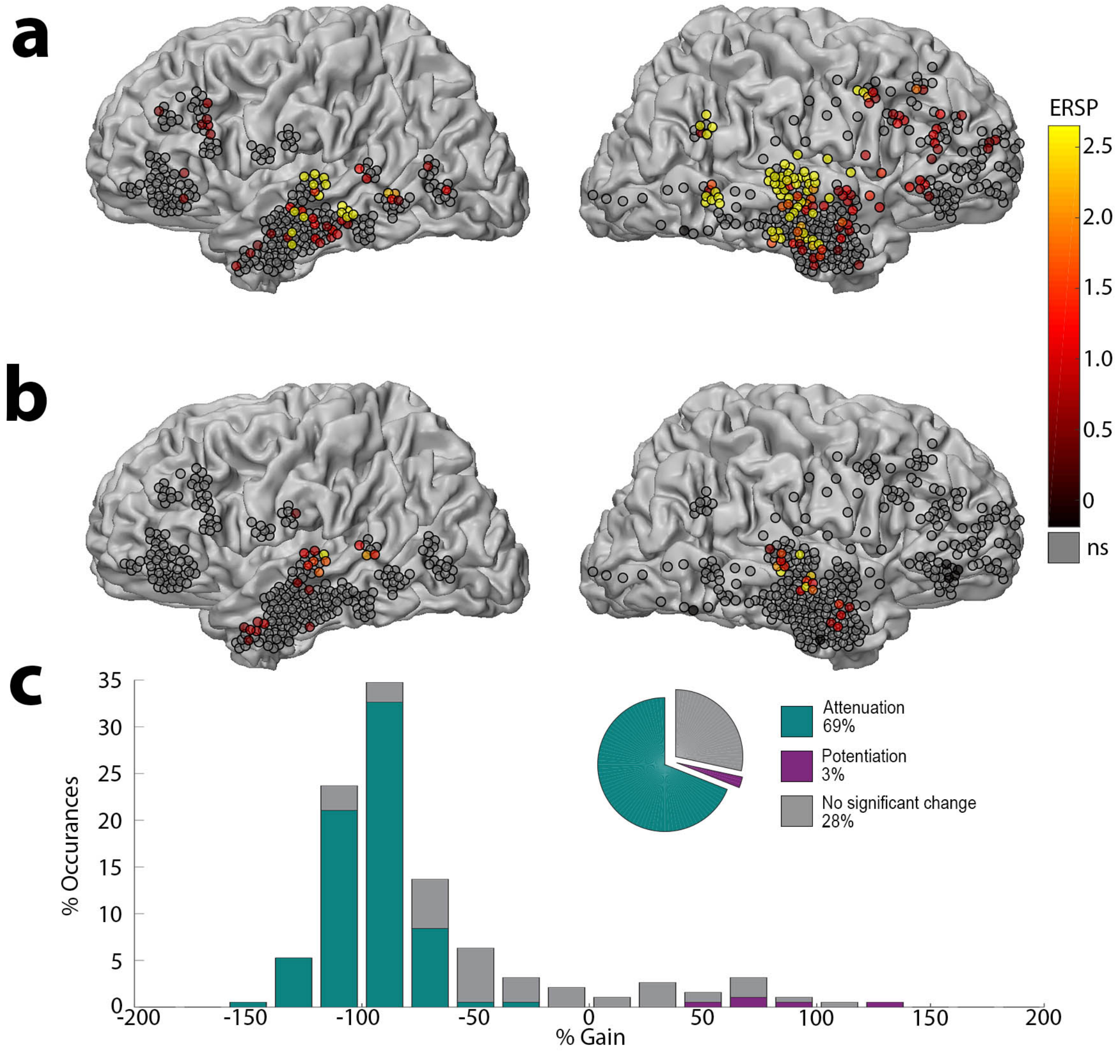


**Fig. S5: Event-Related Spectral Power (ERSP) in iEEG responses to 40Hz click-trains during wakefulness and anesthesia**

(a) ERSP at each iEEG electrode (n = 612 in 9 sessions) in response to 40Hz click-trains during wakefulness, as shown on a grey-white matter boundary surface as seen from lateral view. (b) Same during anesthesia. Note that anesthesia-induced LOC disrupts iEEG responses outside auditory cortex as was seen when quantifying the response via ITPC. (c) Quantification of the 190 iEEG contacts showing significant responses, revealing that 69% of these electrodes undergo significant attenuation under anesthesia, compared to only 3% showing potentiation.

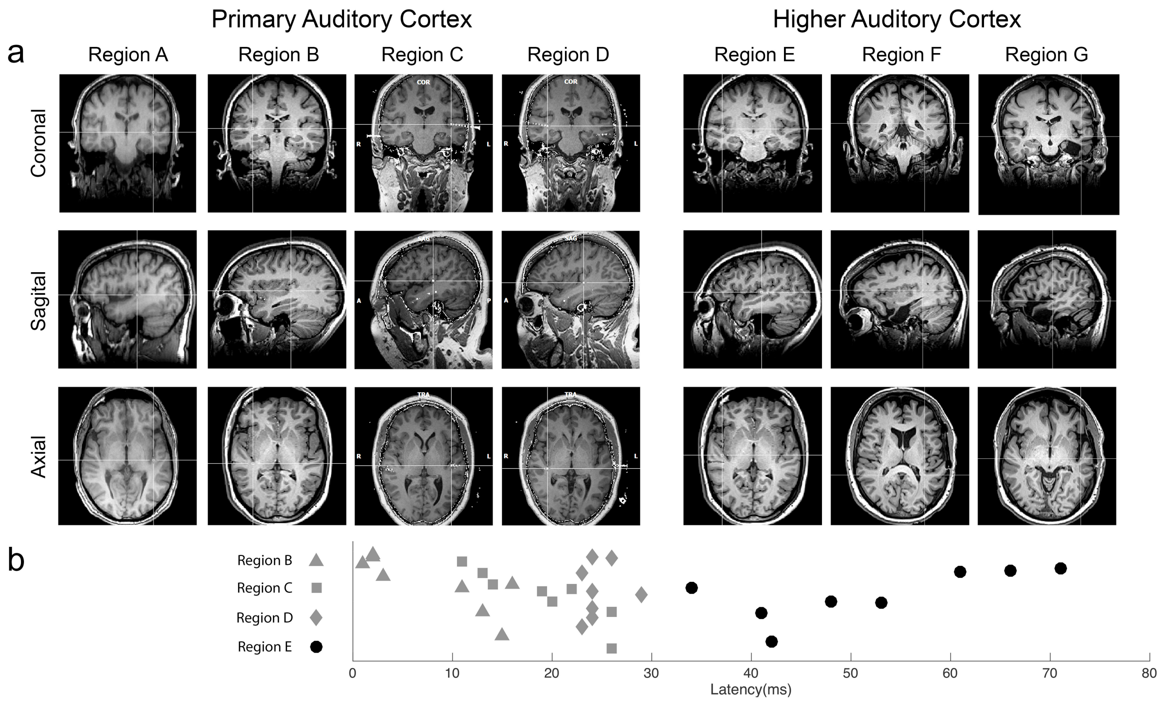


**Fig. S6. Classification of auditory regions to PAC vs. higher-order based on anatomy and response latency.**

(a) Anatomical T1 weighted MRI scans centered on each microwire region, confirming locations within Heschl's gyrus (PAC), or nearby locations (Higher Auditory Cortex). (b) Latency plots for each microwire with sufficient inter-trial coherence to allow latency to be measured. All PAC microwires showed latency < 30ms (mean 17 ± 9 ms), whilst all Region E microwires were > 30ms (mean 52 ± 13 ms).

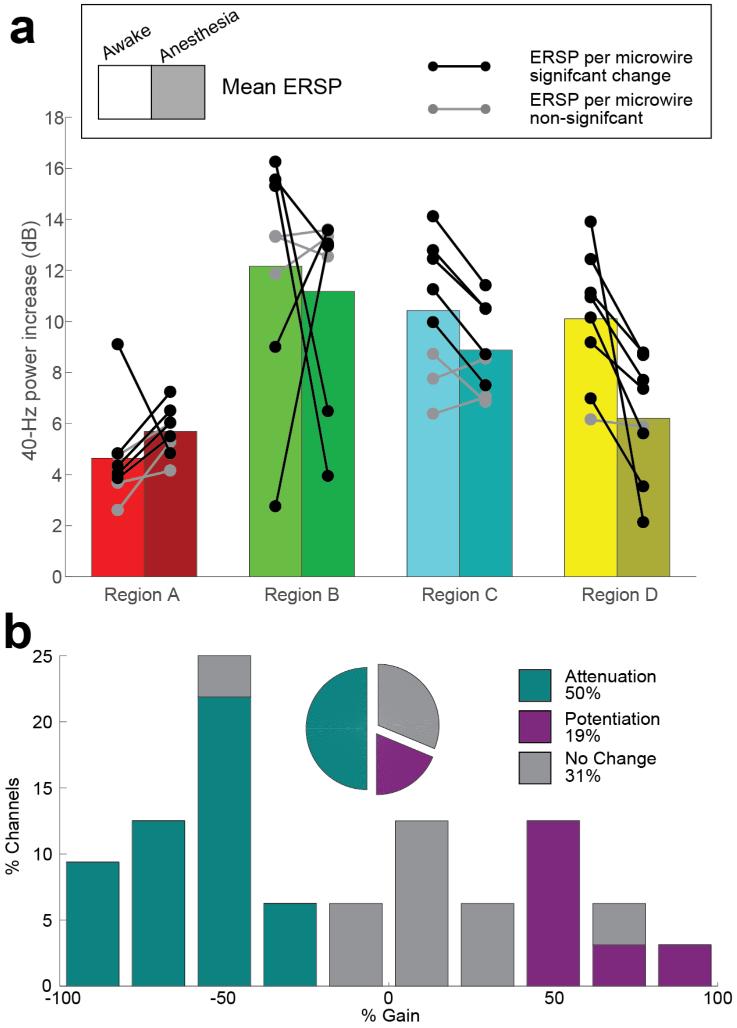


**Fig. S7. Event-Related Spectral Power (ERSP) in primary auditory cortex to 40Hz click-trains during wakefulness and anesthesia**

(a) Event Related Spectral Power (ERSP) at 40Hz in response to click trains across all 32 PAC microwires (circles), and averaged per region (bars), during wakefulness (left bar, bright colors) and anesthesia (right bar, dull colors). PAC microwires showing significant changes ($p<0.05$ Wilcoxon rank sum) are in black, whilst those not showing significant change are marked in grey. (b) Histogram of gain in ERSP under anesthesia in all 32 PAC microwires, divided according to significant attenuation, significant potentiation, and no significant change ($p>0.05$, Wilcoxon rank sum) under anesthesia. As was the case for ITPC (Fig. 3C), LFP responses in PAC are relatively preserved, a profile that is significantly different than observed for iEEG power changes in association cortex (Supplementary Figure 5).

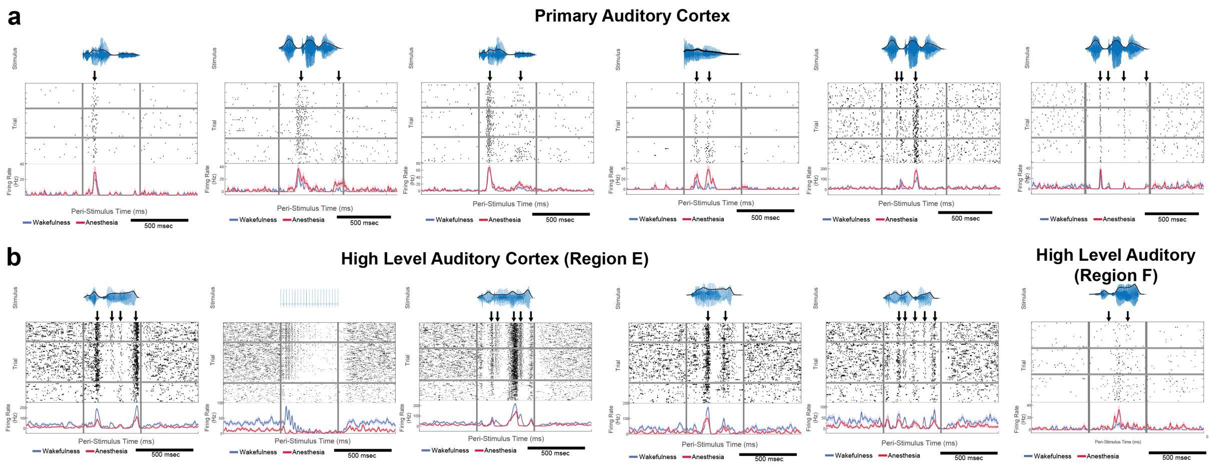


**Fig. S8. Additional examples of neuronal unit spike responses to words upon anesthesia-induced LOC.**

(a) Example raster plots and PSTHs of single-neuron spike responses to words in PAC. Each subpanel shows a different stimulus (click-train or word, waveform on top in blue and its power envelope in black). In each subpanel, rows (top to bottom) mark individual trials during deepening propofol anesthesia. Black arrows indicate identified response components. Blue PSTH time-course, wakefulness; Red PSTH time-course, anesthesia. Vertical gray lines mark stimulus onset and offset. (b) Same as (a) for high-level auditory cortex. Each subpanel shows the response of one neuron to one stimulus and may include one (e.g. in 1st panel) or more (other panels) response components.
